# Activation of the Anaphase Promoting Complex reverses multiple drug resistant cancer

**DOI:** 10.1101/2020.05.26.115337

**Authors:** T.G. Arnason, V. MacDonald-Dickinson, J.F. Davies, L. Lobanova, C. Gaunt, B. Trost, M. Waldner, P. Baldwin, D. Borrowman, H. Marwood, Z.E. Gillespie, F.S. Vizeacoumar, F.J. Vizeacoumar, C.H. Eskiw, A. Kusalik, T.A.A. Harkness

**Author notes:** ***Correspondence to**: Terra G Arnason/Troy A.A. Harkness, **e-mail** or.

## Abstract

Like humans, canines spontaneously develop lymphomas that are treated by chemotherapy cocktails and frequently develop multiple drug resistance (MDR). Their shortened clinical timelines and tumor accessibility make them excellent models to study MDR mechanisms. We previously demonstrated that adjunct treatment of *in vitro* MDR cell lines with insulin-sensitizers effectively restored MDR chemosensitivity and prevented MDR development. This study extends the use of an insulin-sensitizer to clinical and tumor responses *in vivo* in volunteer canines with MDR lymphoma, including assessing changes in MDR protein biomarkers and global gene expression. Longitudinal tumor sampling and analysis of MDR cases throughout treatment allowed a correlation between *in vivo* molecular mechanisms and clinical responsiveness. We found reduced MDR biomarkers within all tumors, yet only one canine entered clinical remission. Analysis of tumor samples during remission and relapse allowed comparison of gene expression profiles. This revealed the Anaphase Promoting Complex (APC), a ubiquitin-E3 ligase regulating cell cycle progression, was impaired during chemoresistance/MDR and restored during remission. Validating *in vitro* tests restored MDR chemosensitivity upon APC activation, supporting the idea that APC activity is an important underlying cellular mechanism associated with treatment resistance, and a novel potential therapeutic target.

## INTRODUCTION

It is estimated that one in two people will develop cancer in their lifetimes, a staggering statistic to consider. Advances in surgical techniques, cancer-specific therapies and small molecule inhibitors have contributed to survival and disease-free duration [1]. However, the development of multiple drug resistance (MDR) to first-line chemotherapeutic agents leaves few, if any, proven treatment options, and may signal a shift to palliative support rather than active therapy [2–6]. MDR can be inherent (prior to treatment) or acquired at any time after initial treatment responsiveness, with recrudescence as much as 20 years later [7,8]. Malignancies of the breast, blood, lung and colon are particularly known for their relatively high rates of treatment resistance and acquired MDR. New treatment options are required for MDR cancers.

Our goals are to identify the underlying cellular pathways involved in MDR development, and to identify novel agents that will promote reversal or prevention of MDR cancers. We have used *in vitro* MDR cell culture models of a variety of human cancers to identify molecular biomarkers and their post-translational modifications (phosphorylation and acetylation) that correlate with both entry to, and exit from, an MDR state [9–13]. Simultaneously, we have evaluated the utility of several insulin-sensitizing oral agents with unexpected anti-cancer activity that returns MDR cells to treatment-sensitive states [10,11,13]. Of relevance here, we found that repurposing the insulin-sensitizing oral drug, metformin, in breast cancer cells *in vitro* proved effective at both reversing preexisting treatment resistance in MDR cell populations, and also preventing MDR onset [13]. Metformin is generally believed to primarily mediate metabolic changes in response to insulin resistance in individuals with Type 2 diabetes through the AMP-dependent protein kinase (AMPK) [14,15]. Although metformin was also shown to slow the growth of multiple human cancer cells *in vitro* [16], its specific effects of MDR malignancy had not been evaluated. We observed that the anti-cancer properties of metformin in MDR breast cancer cells *in vitro* was AMPK-independent and induced histone acetylation via indirect inhibition of histone deacetylases (HDAC) [13]. Compounds with HDAC inhibitory (HDACi) capacity are currently in Phase III cancer trials, with several approved for clinical use [17,18].

In this study we have used the *in vivo* companion canine model of MDR lymphoma with adjunct metformin treatment and observed that MDR protein biomarkers were reduced in all tumors sampled. Microarray analyses of companion canines throughout their treatment protocol, particularly one that entered remission, lead to the discovery that the Anaphase Promoting Complex (APC) was impaired in MDR tumors and that metformin treatment coincided with APC reactivation. Furthermore, we show that *in vitro* activation of the APC by a small chemical APC activator can reverse drug resistance. The APC is a large evolutionarily conserved ubiquitin ligase that is primarily considered to target proteins that inhibit cell cycle progression through mitosis and G1 for ubiquitination and proteasome-dependent degradation [19,20]. More recently, accumulating evidence implicates the APC in cell cycle-independent functions such as neurogenesis, longevity, stress responses and genomic stability [21–24]. The idea that effective APC function is critical to ward off cancer development and/or progression is gaining traction, as the APC has been shown to be important for DNA repair decisions [25,26], and the preservation of genomic stability in humans and yeast [27,28]. APC impairment has also been associated with onset of drug resistance [29,30]. We tested the hypothesis that APC impairment mediates MDR development and demonstrate the APC may serve as a potential therapeutic target that maintains genomic integrity, thereby delaying the development of aggressive treatment-resistant tumors.

## RESULTS

### Companion canines with non-Hodgkin-like lymphoma are strong models of MDR malignancy

Our proximity to the largest veterinary college in western Canada provided us an opportunity to obtain treatment resistant tumor samples representative of *in vivo* events as companion dogs present regularly to the oncology clinic for diagnosis and chemotherapy. Canine lymphoma is orthologous to non-Hodgkin lymphoma of B- and T-cells and has a relapsing-remitting course with a high incidence of terminal MDR transformation [31–33]. The superficial nature of the lymph node enlargements allows for easy, non-invasive tumor sampling. Standard clinical management of lymphoma in companion canines involves repeated weekly appointments, providing an opportunity to sample a given tumor over time during the chemotherapeutic regimen. To test our *in vitro* observations that metformin reverses MDR protein markers and resensitizes MDR cells to chemotherapy, we added oral metformin to the treatment regimes in companion canines that developed MDR lymphoma. We obtained tumor samples from 8 canines in total, 2 from naïve drug sensitive canines and 6 that had failed their drug therapy. Details of the 4 canine subjects used for our complete analyses are shown in Supplemental Table 1. Canine subjects 1-4 were clinically unresponsive to chemotherapy, were identified as having B-cell lymphoma, and all received adjunct oral metformin, as described. Canine subjects 5 and 6 were treatment resistant, but were not treated with metformin and were not followed any further. Canine subjects 7 and 8 were treatment sensitive, were not staged, and did not receive metformin.

### Canines with drug resistant lymphoma overexpress proteins associated with drug resistance, and adjunct metformin therapy results in reduced MDR protein biomarker levels

We used Western analyses to determine the relative abundance of the MDR biomarker, MDR-1, in cancerous lymph nodes as compared to skin control samples, before and after metformin use. MDR-1 is a non-specific drug efflux pump from the ABC trasnsporter family (ABCB1). We used fine needle aspirates (FNAs) to obtain tumor samples from 5 MDR canines (samples 1-4, 6), who all expressed high levels of MDR-1 in their tumor sample compared to their control sample (Fig. 1). We assessed additional MDR protein markers in Canine 3 tumor samples, such as BCRP (Breast Cancer Resistance Protein), HIF1α (Hypoxia Inducible Factor 1 alpha), and phosphorylated S6K, and found them to all be elevated (Fig. 1C). The sample size limited the extent of protein analysis possible. Samples were then taken at the indicated weeks after metformin initiation. Remarkably, MDR-1 protein, and the other MDR protein markers in canine 3, rapidly declined or were undetectable with metformin ingestion (Fig. 1). MDR-1 signal was not detected in canine subjects 7 and 8, as expected, since they were clinically treatment-sensitive (Fig. 1B). Thus, the *in vitro* observations showing loss of MDR markers following metformin treatment [13] are also observed *in vivo*, and potentially clinically relevant as one canine experienced a period of remission following metformin exposure.

**Figure 1.**
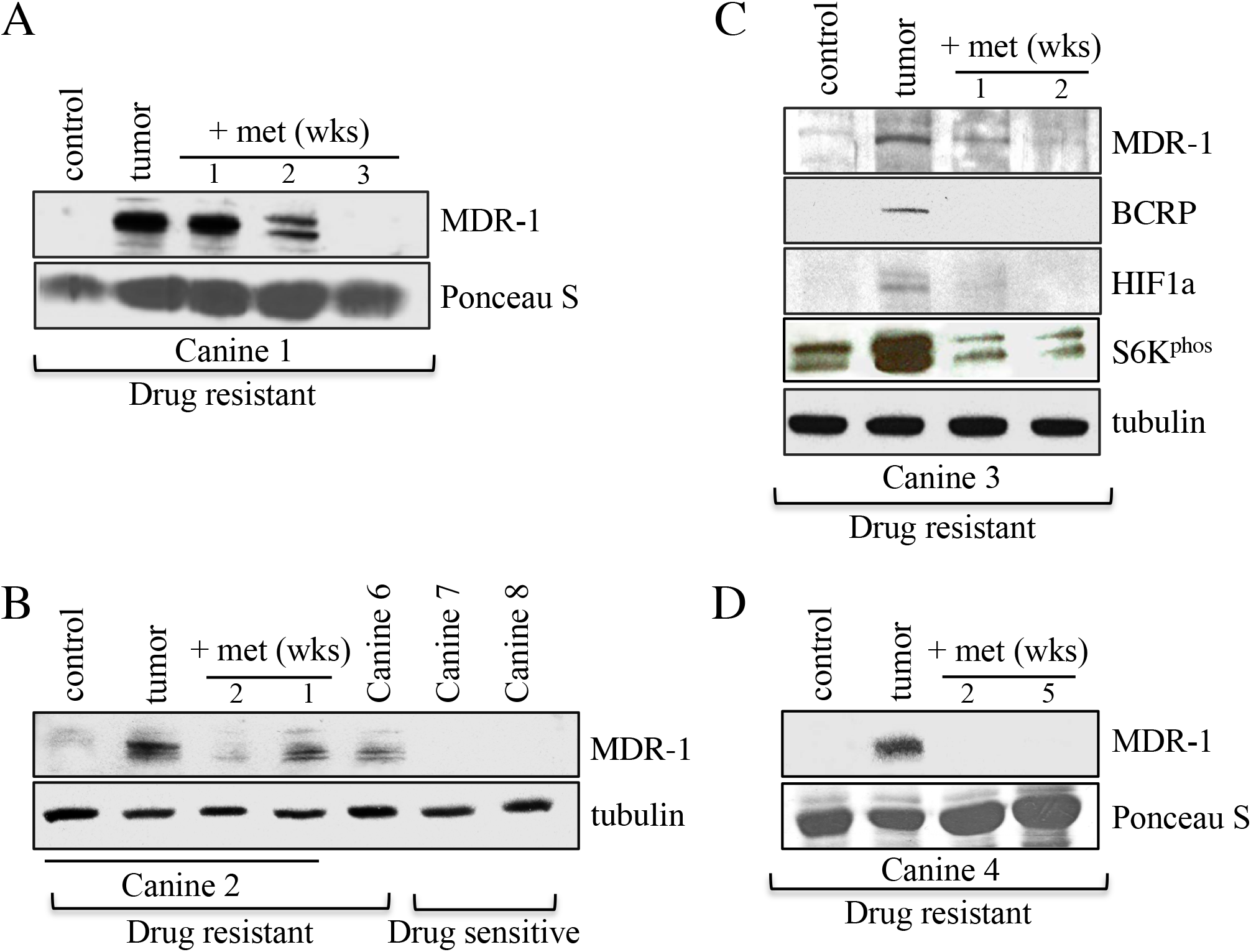
Changes in MDR protein biomarker levels within tumor samples from canine subjects before and after oral metformin addition. **A.** Protein lysates were prepared from Canine 1 that presented with MDR lymphoma. The lysates were analyzed by Western blotting using antibodies against MDR-1. Ponceua S staining was used to ensure equal protein load. The control was derived from skin samples from Canine 2. **B.** Protein lysates were prepared from Canines 2, 6, 7 and 8 for analysis by Western blotting using antibodies against MDR-1 and tubulin. Canines 2 and 6 were recruited with drug resistant lymphoma, while Canines 7 and 8 were drug sensitive. Tumor samples were obtained by fine needle aspirates (FNAs) following 1 and 2 weeks of metformin treatment from canine 2. **C.** Lysates prepared from skin and tumor samples obtained from drug resistant Canine 3, before and after metformin therapy, were analyzed with antibodies against the MDR markers shown, with tubulin serving as the load control. **D.** Protein lysates prepared from skin and tumor samples from drug resistant Canine 4, before and after metformin treatment, were assessed using antibodies against MDR-1, with Ponceau S staining representing relative load controls.

### Metformin as adjunct therapy was nontoxic and well-tolerated

We had already established that the dose of metformin used in this study is nontoxic to canines and generally well tolerated [34,35]. Metformin as monotherapy is not presumed to be highly effective at killing cancerous lymphoma cells, which was supported clinically by the ongoing lymphadenopathy in our subjects. There was a decrease in food interest and this was interpreted to indicate possible gastrointestinal intolerance of metformin [36], a well known side effect in humans. Laboratory investigations spanning metformin monotherapy in these subjects did not show a metabolic acidosis (data not shown), a relevant consideration given that lactic acidosis has been reported as a rare complication of metformin use in humans with renal failure [37,38]. There was no alteration to differential cell counts, with immature bands being equally present before and after weeks of treatment, indicating that rapid cell line expansion was not altered. Blood glucose levels remained normal, as did renal function. The remainder of the bloodwork was unchanged following metformin addition. Although the dual therapy did not, for the most part, result in measurable decreases in lymph node burden, there was a subjective increase in energy and activity reported even with metformin still being administered. A slight elevation in total lactate levels was seen in canine subject 3 after the first week on metformin therapy (3.59 mmol; normal < 2.5 mmol). While metformin is rarely reported to drive lactic acidosis [39,40], the observed rise was not unexpected in rapidly proliferating malignancies such as lymphoma and was likely multifactorial including the comorbid presence of mild anemia and a significant urinary tract infection.

### Microarray analyses of canine tumor mRNA validates increased expression of MDR-1 and reduction of MDR-1 by metformin

We isolated RNA from tumor cells and skin biopsies obtained from 4 treatment resistant companion canines in our study (Canines 1-4). We performed microarray analyses (Agilent Canine Microarrays; 25,000 annotated genes) to identify genes that were differentially expressed in the tumor samples compared to a normal skin sample (Canine 2), and in the tumors before and after metformin treatment (Canines 2 and 4). The datasets have been deposited at the GEO repository (GEO accession # GSE121242; https://www.ncbi.nlm.nih.gov/geo/query/acc.cgi?acc=GSE121242). To validate our Western detection of increased expression of MDR-1, we first analyzed expression of the 40 ABC transporter family members present on the array. We observed expression of *ABCB1* mRNA (encoding MDR-1) above 3 fold change (FC) in only one tumor sample (Canine 4; Supplemental Fig. S1), with mild elevation in canine 2. There were 2 probes for *ABCB1* on the array, and both were elevated in canine 4 well above the 3 FC cutoff (Log base 2 equals a FC of 4), and mildly so in canine 2. *ABCB1* mRNA expression was unaffected in canine 3. Since MDR-1/ABCB1 protein was elevated in all MDR canines, the lack of consistent elevated *ABCB1* mRNA suggests that increased MDR-1 protein levels can occur post-translationally. Only *ABCB3* was elevated above 3 FC in all MDR canines at the mRNA level. Expression was elevated in the MDR canines (with at least three out of four 2-fold or higher) for 4 additional ATP transporters (*ABCA3, ABCC10, ABCG1*, and *ABCA4*). Next, we averaged expression from the tumor samples before metformin treatment from canines 2 and 4, normalized to the control sample, and compared this to the average of canines 2 and 4 treated with metformin normalized to samples before treatment (Supplemental Figure S1). Consistent with metformin effectively reducing the protein levels of ABCB1 (MDR-1; Fig. 1), expression of *ABCB1* mRNA was also reduced by metformin in the 2 canines. Four other ABC transporters showed this pattern, with elevation in the MDR tumor at least 2-fold greater than in the control sample, and reduced by metformin (*ABCA3, ABCC10, ABCG1*, and *ABCB3*). While MDR-1 and other ABC transporters were expressed in these tumors, and while possible that this may be a contributing factor in MDR development, we observed that overexpression of ABC transporters was not a common occurrence in these canines. Thus, we sought to identify other possible mechanisms of MDR development by looking for common differentially expressed genes in the tumors of the 4 MDR canines (canines 1-4).

### A 290 gene set overexpressed >3-fold in tumour versus control was common to the 4 MDR canines

Between 875 (canine 2) and 1365 (canine 3) genes were overexpressed >3 fold in the 4 canine datasets as compared to noncancerous control tissue (Supplemental Table S2). We identified 290 genes overexpressed greater than 3 fold in all 4 canine datasets (Fig. 2A; Supplemental Table S2). Using Cytoscape and Reactome FI [41], we discovered that 146 of these genes were highly associated. In total, 13 different gene clusters were identified (as defined by different colour spheres and asterisks; Fig. 2B) and are listed in Supplemental Table S3. The enriched genes within the network pathways are shown in Supplemental Table S4. A particularly tight interconnected gene cluster (shown in green in Fig. 2B) was identified that was involved in multiple aspects of mitotic regulation. A subsequent connectivity analysis, using the software ‘Search Tool for the Retrieval of Interacting Genes/Proteins’ (STRING, version 10.5), identified 186 genes from the 290 gene set that were involved in multiple interconnected networks/clusters (Supplemental Fig. S2; Supplemental Table S2). One such cluster (Cluster 2) was composed almost exclusively of genes encoding proteins involved in mitotic progression (yellow nodes in Supplemental Fig. S2, highlighted in yellow in Supplemental Table S5). Clusters 3 and 6 also contained many genes involved in mitotic progression, as well as genes required for DNA repair and replication (green and purple nodes in Supplemental Fig. S2; highlighted in green in Supplemental Table S5). Clusters 2 and 4 are joined by the Ras homolog gene family member H (RHOH), while Clusters 2 and 3 are connected by DNA topoisomerase II A (TOP2A). Cluster 6 is interwoven between Clusters 2 and 3. These observations demonstrate that dysregulation of a tightly connected network of genes involved in cell cycle progression and genome integrity may drive MDR development.

**Figure 2.**
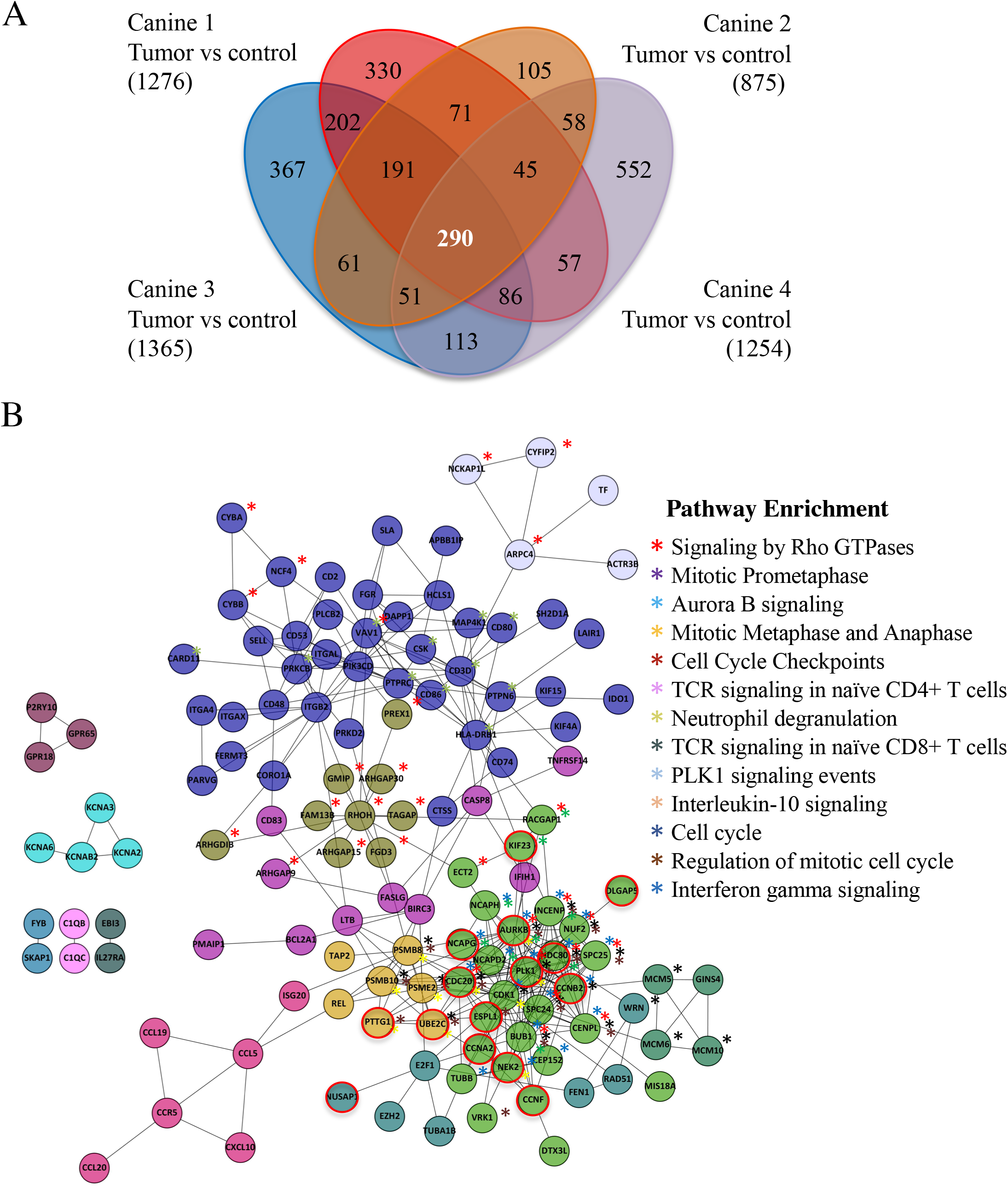
Canines with MDR lymphoma tumors express a common set of highly interactive genes. **A.** A Venn diagram highlights a set of 290 genes overexpressed in the tumors of the four MDR canines studied. Agilent canine arrays containing over ~25,000 annotated canine genes were used to analyze mRNA obtained from tumor samples in four MDR subjects (Canines 1-4) and normalized to control skin tissue mRNAs (Canine 2). The fold change (FC) was determined by comparing the Log base 2 expression levels from the tumors with the control. A Log base 2 of 1 is equivalent to a FC of 2. A FC >3 was used as a cut-off. Bracketed numbers reflect the total number of genes above the FC 3 threshold in that subject. **B.** Cytoscape was used to analyze the 290 gene set. 146 genes were found to form 13 highly interconnected nodes. Network pathways found to be enriched within these nodes are shown. Astericks denoting genes within the nodes that are enriched within network pathways are color coded. Nodes circled in red define Anaphase Promoting Complex substrates.

Both Cytoscape with ReactomeFIViz and STRING identified the same cluster of mitotic genes, in which many of the genes encoded substrates of the Anaphase Promoting Complex (APC; 20 genes, marked by *, Supplemental Table S5; nodes circled in red in Fig. 2B). The APC is a ubiquitin E3 protein ligase that targets proteins that inhibit cell cycle progression for ubiquitin- and proteasome-dependent degradation [20,42]. APC regulatory and substrate mRNAs often peak during G2/M, then decline once the protein is degraded [43,44]. Thus, if APC substrates are elevated, even at the mRNA level, then this suggests APC function is impaired. Furthermore, proteins associated with the Spindle Assembly Checkpoint (SAC) and the kinetochore, which predominantly inhibit APC activity prior to mitosis [45], were elevated (15 genes, marked by **, Supplemental Table S5). Moreover, proteins that maintain chromosome condensation (the condensin complex) were also elevated (3 genes; marked by ***, Supplemental Table S5). Interestingly, a recent study comparing differentially expressed genes between normal human ovarian tissue and ovarian cancer, and carboplatin-sensitive and resistant ovarian tumors [46], identified a similar set of genes identified in our canine study; of the 5 top genes identified in their study (GNAI1, NCAPH, MMP9, AURKA and EZH2), three were identified as elevated in our study (NCAPH, AURKB (rather than AURKA) and EZH2). Taken together, our work suggests that MDR development in these 4 canines is associated with impaired APC activity. Consistent with this, we demonstrated using the Cancer Genome Atlas (TCGA) (https://portal.gdc.cancer.gov/) database that the APC substrates Securin, HURP [24] and Cyclin B1 (Supplemental Fig. S3) are elevated in at least 24 different human cancers from patient samples.

### Mitotic and G1 APC substrate gene expression is elevated in treatment resistant tumors compared to normal controls

The APC targets specific proteins for degradation in both mitosis (M) and G1, and we asked if there was a bias within MDR samples compared to control for target expression in the array. Different coactivators interact with the APC to promote cell cycle progression; CDC20 during mitosis (APC^CDC20^) and CDH1/FZR1 to exit mitosis and progress through G1 (referred to as APC^CDH1^ hereafter) [19–21]. Therefore, we separated all known APC targets present in the array into M vs G1 targets (11 mitotic substrates and 21 G1 substrates) to assess if there were differences in expression between the clusters (Fig. 3). We consistently observed that the mitotic targets were overexpressed in MDR tissue on average to a greater extent than that observed for G1 substrates (Supplemental Fig. S4), although many G1 substrates were also highly expressed. In some cases APC^CDH1^ will target residual APC^CDC20^ substrates causing an overlap (PLK1, Securin and Cyclin A2 [21]). This analysis revealed unique differences between APC target gene expression in transcripts within all 4 MDR samples.

**Figure 3.**
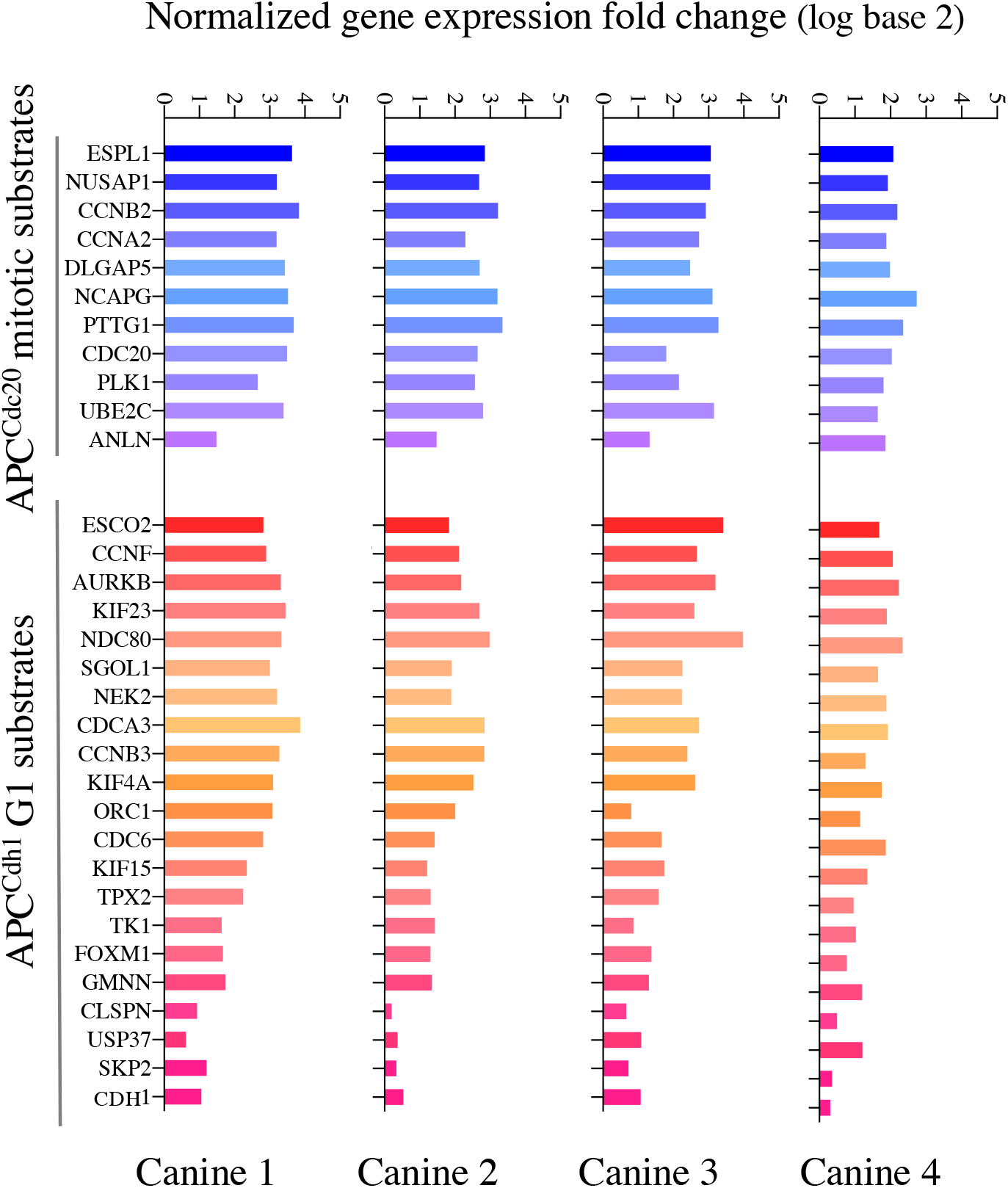
mRNAs encoding APC substrates are elevated in the 4 MDR canines. All known APC mitotic and G1 specific substrates, to the best of our knowledge, that were present in the canine microarray were assessed for differential gene expression in MDR tumor samples compared to control skin samples. The histogram in shades of blue reflects APC^Cdc20^ mitotic substrates, while those in shades of red define APC^Cdh1^ targeted G1 substrates. Samples from the 4 MDR canines are shown individually. Average values are shown in Supplemental Figure 4.

The coordinated gene expression of functionally associated genes may be due to activation by one or more common transcription factors for APC substrates. To gain insight into this possibility, we implemented a Cis-element Over-representation (CLOVER) Analysis [47] using the TRANSFAC database to analyze the 290 gene list. This analysis identifies over-represented transcription factor binding sites within the promoters of genes. This analysis revealed that 230/290 of the genes have FOXO3A and/or FOXM1 binding sites, as well as 241 sites for the SP1-4 transcription factor family. A Venn diagram shows that 225/290 of the genes contain sites for all 3 sets of transcription factors (Supplemental Fig. S5). Our finding that FOXM1 may be highly involved in the expression of the 290 gene list is significant since FOXM1 is a known APC^CDH1^ substrate [48]. As can be seen in Fig. 3, FOXM1 expression was elevated in all canine tumors, providing a mechanism for how the APC substrates can be coordinately overexpressed in the canine tumors. Further work will need to be done to clarify this observation.

### Microarray reveals reversible changes in APC target gene expression that correlate with altered clinical treatment responses

Canine 4 exhibited a successful remission (temporary) based on the near disappearance of all palpable lymph nodes. It should be highlighted that this dog had failed both repeat CHOP therapy and rescue therapy in the months prior to our study entrance. Clinical remission occurred when CHOP was again trialed but with the addition of oral metformin. After 8 weeks in remission the subject relapsed and expired. RNA samples were obtained from the same lymph node tumor location prior to metformin therapy, during remission while on metformin therapy as the tumors shrank, and from the tumor after relapse. This self-controlled analysis identified a set of 27 genes that were upregulated >3 FC in the MDR tumor, decreased >2 FC upon remission and then elevated once again >3 FC upon relapse (Fig. 4A) as compared to the skin control. We performed STRING on this set of genes and found that they were highly enriched in APC substrates that were overexpressed at a time of clinical unresponsiveness to therapy (Fig. 4B, highlighted with a black ring around the node; Supplemental Table S6). This further supports the concept that APC function is correlated with clinical disease responsiveness. Further analyses of APC mitotic and G1 substrates in our three temporal samples from canine 4 revealed that the mitotic APC substrates that were elevated in tumors were remarkably reduced during remission, and upon relapse, were elevated beyond the initial MDR lymphoma cells (Fig. 4C). This was also observed for the bulk of the APC G1 substrates (Fig. 4C). We validated the gene expression changes detected through the microarray analysis by performing qRT-PCR on the original FNA samples from chemoresistant and remission samples in Canine 4, for the genes encoding HURP (*DLGAP5*) and Securin (*PTTG1*; Fig. 4D). We confirmed that gene expression is significantly decreased in the treatment-sensitive/remission sample as compared to the MDR sample.

**Figure 4.**
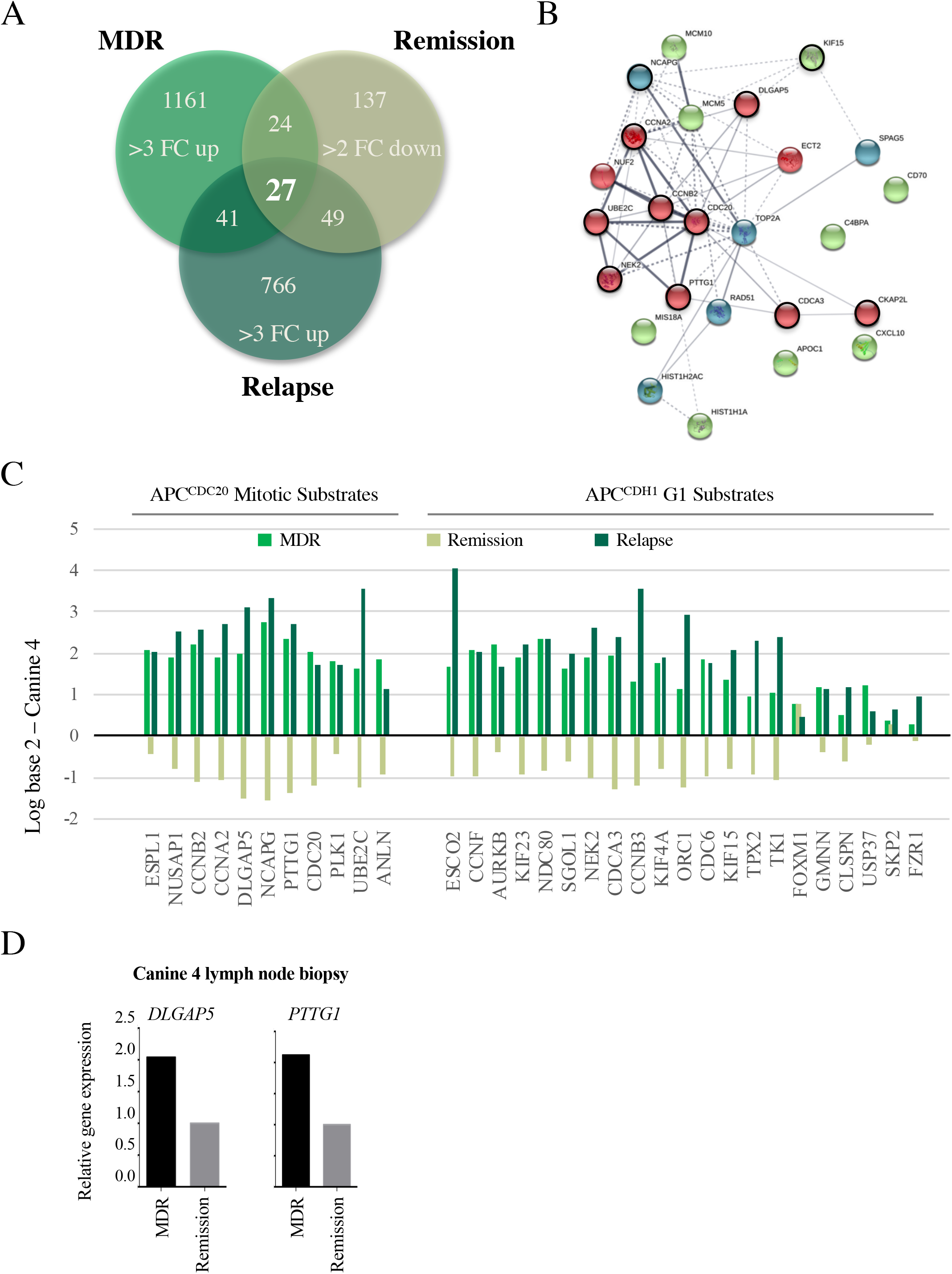
Changes in APC target gene expression correlate with altered clinical responses. **A.** Lymph node FNA samples were obtained from Canine 4 before metformin treatment when the tumor was chemoresistant (MDR), during remission following metformin treatment (Remission), and then after loss of remission (Relapse), with differential gene expression compared to normal skin samples using canine microarray. All Canine 4 samples were taken from the same cervical lymph node. A Venn diagram was used to determine similarities in genes overexpressed 3 FC in MDR tumors, 2 FC down-regulated during remission, then 3 FC upregulated following relapse. This is predicted to identify genes specifically involved in the MDR phenotype that correlate with clinical responsiveness. From this analysis, 27 genes were found to be overexpressed in chemoresistant cells, reduced upon remission, then overexpressed once again when remission failed. **B.** A STRING analysis indicates that the 27 genes were highly interconnected, with the majority of the genes encoding APC substrates (red nodes, circled in black), and genes encoding proteins required for chromosome maintenance (green and blue nodes, APC substrates circled in black). **C.** Differential gene expression was determined for the APC substrates present in the array (Figure 3) compared to control skin samples, as correlated with clinical response to treatment at the time of sampling: MDR, remission, and relapse. **D.** Microarray results were validated by qRT-PCR of original FNA aspirate samples for *DLGAP5* (encoding HURP) and *PTTG1* (encoding Securin) in Canine 4 at study entry and remission.

### APC substrates are elevated *in vitro* in OSW lymphoma cells selected for DOX resistance

We queried whether APC impairment may be a common mechanism in MDR transformation. To extend the observations that APC activity is impaired *in vivo* in drug resistant clinical samples, we obtained the canine OSW lymphoma cell line [49] to test whether our *in vivo* observations were valid *in vitro*. We selected OSW cells for resistance to Doxorubicin (OSW^DOX^) according to our established methods [12,13]. The MDR status of the OSW^DOX^ cells was confirmed by the overexpression of protein biomarkers of MDR, including BCRP, MDR-1, and HIF1α (Fig. 5A), and the detection of significant resistance to DOX cytotoxicity (Figure 5G). Quantitative PCR (qPCR) of OSW parental (OSW^sens^) and OSW^DOX^ cells confirmed that APC substrate genes *PTTG1, DLGAP5* and the MDR marker *MDR1* were elevated at the mRNA level in *in vitro* selected cells (Figure 5B), paralleling the *in vivo* MDR canine tumor samples. As expected with a defect in APC activity, we observed that APC protein substrates were increased specifically in the MDR cell populations, as noted by the increased protein abundance of HURP, Cyclin B1 and Securin (Figs. 5C and 5D).

**Figure 5.**
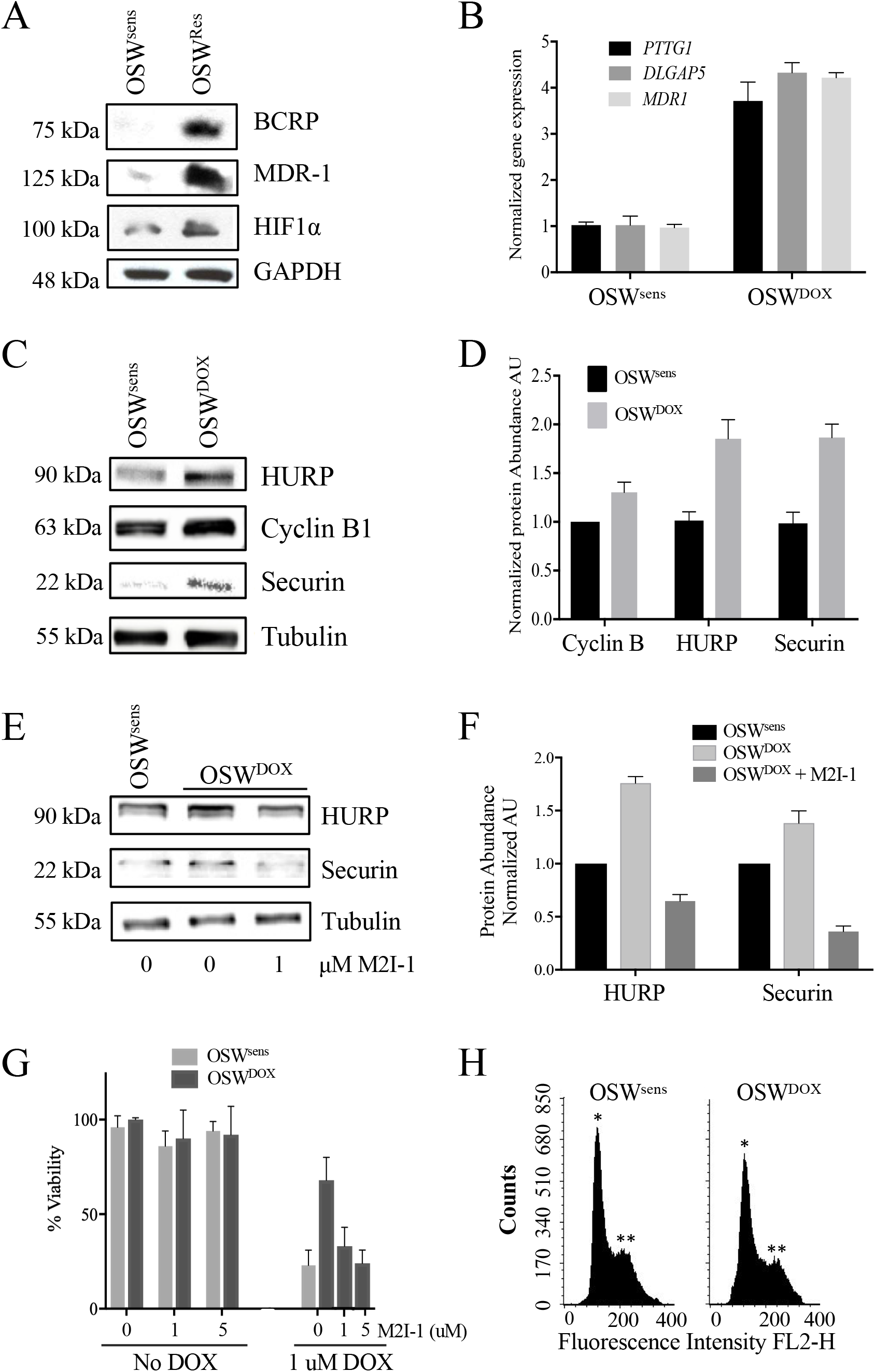
Reversible APC dysfunction in MDR cell populations is validated to occur *in vitro* and correlates with chemosensitivity. **A.** OSW canine lymphoma cells (OSW^sens^) were selected for resistance to DOX (OSW^DOX^). Protein lysates from chemosensitive and resistant populations were analyzed by Western analysis for multiple MDR biomarkers. **B.** qRT-PCR analysis of APC substrate and MDR marker genes in matched populations (n=3). **C.** OSW cell lysates from (**A**) were tested for APC-target protein abundance. **D.** Quantification of Western protein abundance using ImageJ, normalized to Tubulin levels (3 rpts). **E.** OSW canine lymphoma cells selected for DOX (OSW^DOX^) were exposed to the APC activotor M2I-1 (1 μM) for 18 hours or left untreated. Untreated parental cells (OSW^sens^) were used as a comparison. Protein lysates from these cells were analyzed by Westerns for the abundance of APC protein targets. **F.** Protein bands in **E** were quantified using ImageJ, normalized to Tubulin levels, and plotted. Three separate Westerns were analyzed. **G.** OSW^sens^ and OSW^DOX^ cells were pretreated with 1 or 5 μM M2I-1 for 18 hours, then exposed to 1 μM Doxorubicin for 48 hours. Cell viability was measured using Trypan Blue. Three biological replicates were performed. **H.** Flow cytometry of sensitive and resistant OSW asynchronous cell populations was performed to determine distribution of cell cycle phases. Propidium iodide fluorescence was measured to detect DNA content (*, n=1; **, n=2).

### Activation of the APC reduces APC substrates in MDR cells and resensitizes MDR cells to therapy

If impaired APC activity correlates with MDR development, we predicted that activating the APC should reverse the MDR phenotype by resensitizing these cells to chemotherapy. To directly investigate this idea, we obtained a commercially available APC activator called MAD2 Inhibitor-1 (M2I-1) [50]. This small molecule compound disrupts the MAD2-CDC20 interaction, thereby negating SAC inhibition of the APC. This results in the release of CDC20 to enable interactions with, and activation of, the APC. We treated OSW^DOX^ cells with 1 μM M2I-1 for 18 hours (previously optimized, data not shown) and assessed the protein levels of the APC substrates HURP and Securin, and the load control tubulin (Fig. 5E). As shown in the quantitative analysis in Fig. 5F, HURP and Securin levels in OSW^DOX^ cells dropped to levels lower than in OSW^sesns^ cells upon M2I-1 treatment. This is consistent with the idea that APC E3 activity was being restored with the M2I-1 compound. To test if APC activation correlated with a simultaneous enhancement in chemosensitivity, we tested the effects of M2I-1 on cell viability in the presence of a toxic dose of DOX (1 μM) in OSW^sens^ and OSW^DOX^ cell populations (Fig. 5G). M2I-1 (1 and 5 μM) proved to be nontoxic to both parental and selected cells in the absence of DOX. However, there was a striking dose-dependent killing of OSW^DOX^ cells when M2I-1 and DOX were used in combination that reached the chemosensitivity of OSW^sens^ cells, indicating reversal of resistance. This is consistent with a recent study showing that M2I-1 disrupts CDC20-MAD2 interactions *in vivo*, leading to increased sensitivity of cancer cell lines to nocodazole and taxol *in vitro* [51]. Since it appears that the APC is compromised in OSW^DOX^ cells, which should result in impaired progression through mitosis, we expected that MDR cell populations may accumulate in M-phase with a 2n compliment of DNA. However, this was clearly not the case, as flow cytometry showed that both OSW^sens^ and OSW^DOX^ cell populations are predominatly in G1 (Fig. 5H). It was previously noted that tumor cells with elevated levels of APC substrates, but that are in G1, have accelerated cell cycle progression, and are associated with a higher risk of relapse, with high-grade triple negative breast cancer subtypes [52]. Taken together, the experiments presented here support our hypothesis that APC activity is impaired in MDR cells, and that chemical activation of the APC in MDR cells resensitizes MDR cells to chemotherapy.

## DISCUSSION

In the work presented here we aimed to elucidate the *in vivo* molecular mechanisms involved in the progression of cancer to an MDR state and the molecular networks involved in reversing MDR. Our previous *in vitro* work demonstrated that the insulin sensitizing medication metformin could restore the chemosensitivity of human MDR breast cancer [13]. Furthermore, pretreatment of MCF7 with metformin delayed selection against DOX, indicating a prevention of MDR development. This may in part explain the meta-analysis results indicating that individuals taking metformin for Type 2 diabetes therapy not only have a lower incidence rate of many common cancer types, but also appear to respond more robustly to therapy [53–55]. Translation of our *in vitro* results to an *in vivo* model could provide enormous potential for humans who develop resistance to therapies designed to treat their disease, and reveal the underlying mechanisms that are altered specifically in MDR malignancies.

The canine model of cancer is considered an excellent model for human translation [56–59] and was used in our study to assess the effects of metformin on canines with MDR lymphoma. Six canines that presented to the Western College of Veterinary Medicine (WCVM) at the University of Saskatchewan with treatment resistant lymphoma were recruited into our study. They were all found to overexpress MDR-1, an ABC drug efflux transporter found elevated in many cases of drug resistant cancers in humans [9,60], specifically in their tumors. In the 4 cases tested, metformin reduced MDR-1 protein levels, similar to what we observed *in vitro* [13]. One canine (Canine 4), after failing multiple treatment rounds and all previous rescue therapies, did go into remission, providing evidence that metformin has the potential to reverse drug resistance *in vivo*. However, it should be noted that the canines enrolled in our study were terminally ill, and although there were signs of improvement in their behavior, there was little sign of reduced tumor size aside from Canine 4. Future studies will focus on naïve dogs as they enter the clinic for a blinded study with metformin to determine whether metformin can delay the onset of drug resistant lymphoma.

We obtained tumor RNA from 4 canines that was used to determine differential gene expression using canine microarrays in tumors compared to skin controls. We also obtained tumor RNA from 2 canines (Canines 2 and 4) treated with metformin, and from the one canine (Canine 4) that went into remission and later relapsed. The tumor samples were obtained from the same tumor over time, allowing a direct comparison of gene expression as tumor behavior changed. Since our initial assessment asking whether ABC transporter genes are elevated in MDR tumors did not reveal a general role for ABC transporters in MDR development (Supplemental Figure S1), we looked for common genes overexpressed in all 4 canine tumor samples prior to metformin treatment. Our analyses of overexpressed genes in the MDR tumors of the 4 canines revealed a common set composed of 290 genes overexpressed at least 3 fold as compared to non-cancerous control tissues (Fig. 2A). We used Cytoscape and STRING databases to analyze this gene set and discovered that 146 and 186 of the 290 genes, respectively, were highly associated (Supplemental Fig. S2; Supplemental Tables S2, S3 and S5). These genes are highly enriched for those encoding proteins involved in mitotic cell cycle progression, especially degradation substrates of the APC and components of the kinetochore and SAC. Other genes that encode signaling molecules, and proteins required for DNA-dependent functions (ie., DNA repair, chromosome condensation and kinetochore/centromere assembly) were also enriched and are important for progression through mitosis. These observations clearly indicate that tumor development is multifactorial, and includes an impaired progression through mitosis, potentially controlled by APC activity.

To validate the idea that APC is impaired in canines with MDR lymphoma, resulting in the likely accumulation of APC substrate mRNAs, we determined the fold change of the APC substrates we could identify in the canine array (Fig. 3). Almost all of the 32 substrates identified were elevated at least 3 FC, including all mitotic substrates in all 4 canines and most of the G1 substrates (Fig. 3). Previous work has also demonstrated multiple APC substrates to be elevated in various cancer cell lines [61–63]. Furthermore, recent work comparing differential gene expression in drug sensitive and resistant ovarian tumor datasets [46] revealed a similar collection of genes to those identified by us; half of the 30 highly connected nodes they identified were associated with APC function or substrates. Consistent with these observations, remission and relapse of Canine 4 following metformin treatment identified 27 genes that were upregulated in the tumor, down-regulated following metformin treatment, then upregulated again following relapse (Fig. 4A). These 27 genes were highly interconnected and mostly composed of APC substrates (Fig. 4B), clearly placing the APC at a pivot point as a critical activity in maintaining cell health.

Elevated APC substrates can arise from multiple mechanisms, including failure to be degraded due to dysfunctional APC E3 ligase actiity, but also due to dysregulation of their gene expression. The APC targets proteins for degradation, not mRNA, but APC substrates are often transcriptionally active just prior to when they are required [43,44]. This is controlled by transcripton factors that are themselves degraded during the cell cycle by the APC, including FOXM1, which was found to have promoter recognition sites in many known APC target genes (discussed in [48]). The mRNAs encoding many APC substrates in yeast are also cell cycle regulated, with synthesis of their corresponding proteins peaking following the mRNA expression peak [64–66]. In fact, transcription factors that transcribe yeast APC substrates, such as Fkh1 and Ndd1 are targeted by the APC [27,67], thereby creating feedforward loops. This loop is further layered by the observation that the yeast Fkh2 transcription factor (also a likely APC target; our unpublished observation) recruits Ndd1 to chromatin of additional G2/M specific APC subunit promoters [68]. The accumulation of APC substrate mRNA, at least in part, is a result of impaired APC-degradation of the transcription factors responsible for their synthesis.

Elevated APC substrates can have many biological consequences that are relevant to cancer behavoir. First, many substrates, such as CCNB2, PLK1 and CDC20, are both APC activators and APC substrates. Elevation of these mRNAs and proteins could result in persistent APC activity that impacts the cell. Increased accumulation of APC substrates such as CDC20 has lead to suggestions that inhibition of the APC, via down-regulation of CDC20, is important for cancer treatment [61,62]. However, other proteins, such as Securin and HURP are not APC activators, but merely substrates. Elevation of these substrates, and many other APC substrates are linked with cancer and other diseases [24,61,63], and this is consistent with the idea that impaired APC activity is a primary driver of tumor progression [25–30]. Although many APC substrates are elevated in this study, suggesting a global fault in APC activity, genes encoding SAC components and the kinetochore (BUB1, CENPL, CEP152, NUF2, SPC24, SPC25, NEK2, INCENP, SGO1L, MLF1IP, CASC5, NDC80, and ASPM), which inhibit the APC prior to anaphase [45,46,69–71], are also elevated (Supplemental Tables S2 and S3). Elevated expression of SAC and kinetochore components (NCAPG, NCAPG, SKA3, AURKB, NEK2) are associated with cancer [72–77], and is consistent with dysregulated upstream APC activation.

To further validate our results that APC activity is impaired in MDR cells, RNA isolated from Canine 4 tumor tissue before and after entry to remission confirmed that the gene expression of APC substrates HURP and Securin were indeed reduced with clinical improvement (Fig. 4D). Validation *in vitro* using canine OSW lymphoma cells selected for resistance to DOX confirmed that APC substrates were elevated in MDR cells compared to parental chemosensitive cells (Figs. 5B-5D). We further proposed that activation of the APC may increase cell health and may resensitize MDR populations to chemotherapy. Activation of the APC using a commercially available small compound that inhibits the MAD2-CDC20 interaction, M2I-1 [50], reduced APC substrate levels in the selected OSW cells and resensitized the selected cells to DOX again (Figs. 5E-5G). Our results demonstrate that impaired APC function may be a critical trigger leading to progression towards aggressive tumor behavior and decreased treatment responsiveness.

We observed that OSW^DOX^ cells are primarily in G1 with elevated levels of APC mitotic substrates (Fig. 5H). A previous study of 182 breast cancer patient samples revealed that 58% of cases were characterized contained elevated levels of the mitotic APC substrates GMNN, PLK1 and Aurora A yet stained abundantly for G1/S markers [52]. These samples were from patients with high grade triple negative breast cancers and at high risk of relapse. Cells with elevated mitotic markers but in G1 can occur by mitotic slippage, where cells can bypass a mitotic arrest and progress into G1 [78]. Mitotic slippage can occur when the SAC is overridden and this is associated with an MDR phenotype [78]. Our work revealing that OSW^DOX^ cells exhibit high levels of APC mitotic substrates in a population predominantly in G1 is consistent with a higher grade cancer and has the potential to serve as a diagnostic marker for aggressive tumor progression. We propose that targeting the APC for activation through pharmaceuticals may reverse MDR behavior, and thereby provide much needed treatment options for those suffering from untreatable malignancies.

Taken together, the results presented here validate the companion canine as a powerful and dynamic model of MDR cancer that has allowed us to determine that the APC is important in cancer behavior and treatment sensitivity to chemotherapy. Impaired APC activity correlates with poor clinical outcomes, as well as MDR behavior. Exogenous activation of the APC corrected the impairment and restored chemosensitivity *in vitro*. Our work provides insight into the APC as a novel therapeutic target that may offer hope for individuals presenting with treatment-resistant malignancies, and may prove useful in preventing the development of resistance in cancer patients in the future.

## MATERIALS AND METHODS

### Cell lines and materials

Canine OSW lymphoma cells were obtained from American Type Culture Collection (ATCC) in Manassas, VA, USA. Cells were cultured in 100 mm tissue culture dishes (Nalgene) in a humidified atmosphere (5% CO2) at 37°C containing RPMI 1640 media (Hyclone) with 10% FBS and antibiotics. All treatment compounds were reconstituted in dimethylsulfoxide (DMSO) except metformin (Sigma) which was reconstituted in molecular-grade water and filter sterilized prior to use. Drug treatments were applied at the concentrations and times as indicated. Flow cytometry was performed as described previously [10].

### Doxorubicin-selection of MDR cell lines

OSW parental cells were selected for drug resistance using previously described protocols [12] with initial selection in the presence of 1 uM Doxorubicin (DOX) for 48 hours. Following this treatment, the cells were washed with sterile PBS and allowed a 3-day recovery period. Drug resistance selection pressure was then reapplied to the cells by subculturing in the presence of 100 nM DOX for 2 weeks with fresh media changes every 3 days. Following the selection period the cells were subjected to MDR-1 Western blot analysis to verify the establishment of drug resistance and sensitivity to DOX using an MTT assay.

### Western blot analysis

Non-adherent OSW cells were captured by centrifugation at 1000 rpm and washed once with sterile PBS. Cell pellets were transferred to microtubes and resuspended in ice cold RIPA buffer. The cell suspensions were pulse sonicated and centrifuged to remove cell debris. The resulting cell lysates were subjected to Bradford protein analysis (BioRad) to facilitate equivalent protein loading during Western analysis. Cell lysates were treated with 5X Laemmli buffer containing ß-mercaptoethanol, boiled to reduce viscosity and resolved by sodium dodecyl sulphate polyacrylamide gel electrophoresis (SDS-PAGE), and transblotted onto nitrocellulose membranes. Transblot efficiency was verified prior to immunoprobing by nonspecific Ponceau S protein staining of the membranes. Following primary antibody incubation overnight at 4°C, the blots were probed with a 1:10000 dilution of a secondary horseradish peroxidase (HRP) secondary antibody. Primary antibodies against MDR-1 (Abcam, 1:500, 180 kDa), BCRP (Abcam, 1:1000, ~70 kDa), HIF1α (Santa Cruz Biotechnology (SCBT), 1:500, 110 kDa), S6K^phos^ (SCBT, 1:1000, 60 kDa), HURP (Abcam, 1:1000, 120 kDa), Cyclin B1 (Sigma, 1:1000, 70 kD), Securin (Abcam, 1:1000, 29 kDa), GAPDH (Abcam, 1:1000, 55 kDa) and tubulin (Sigma, 1:1000, ~50 kDa), were used in this study. The antigen/antibody target signals were detected using an enhanced chemiluminescent detection kit (ECL-BioRad) and chemoluminescent detection using BioRad VersaDoc molecular imager and Software. Cells were viewed with an Olympus BX51 fluorescence microscope 100x objective equipped with an Infinity 3-1 UM camera. Images were collected using Infinity Analyse software version 5.0. A.

### MTT assay

MTT (3-(4,5-dimethylthiazol-2-yl) 2, diphenyl-tetrazolium bromide) was used to measure the antiproliferative effects of the compounds used in this study. MTT is a colorometric cell proliferation assay based on the reduction of the yellow MTT compound to a purple-colored formazan product in the mitochondria of viable, living eukaryotic cells. Cancer cells were cultured in 6 well multi-well plates in phenol red-free medium to avoid interference with the analysis of the purple formazan product. The formazan product and spectrophotometric analysis was performed at 570 nm as previously described [12].

### Companion canine recruitment and characteristics

During routinely scheduled clinic visits at the Western College of Veterinary Medicine (WCVM), study candidates were identified and consent obtained. The 8 canines in the study had been previously diagnosed with lymphoma and underwent initially successful chemotherapy. Six of the canines recruited presented to the WCVM with relapse and generalized lymph node enlargement. The canines had at least 2 previous treatment failures and were recruited into the study upon spontaneous relapse and positive MDR biomarkers (MDR-1). Inclusion criteria included the need for owner consent; their ability to financially contribute to the treatment cost, time and travel required to receive chemotherapy; owner willingness to administer oral metformin tablets up to twice daily; and a life-expectancy of at least 6 weeks. The study protocol was approved after a full review by the University Animal Care Committee - Animal Research Ethics Board at the University of Saskatchewan (AREB# 20120063). Upon recruitment by one of two veterinary oncologists at the WCVM, the canines underwent a complete physical exam and staging that included bloodwork (complete cell counts, liver panel, electrolytes, and renal function testing), thoracic radiographs, and abdominal ultrasound prior to initiation of chemotherapy, the choice of which was left to the discretion of the treating clinicians. Oral metformin tablets were dosed on body weight with a maximum dose of 10 mg/kg, initiated as once-daily and increased as tolerated to twice daily at weekly clinic appointments.

### Canine clinical assessment and sample retrieval

Upon recruitment, each companion canine was examined, and samples were taken both for staging and our analyses. At the first visit, a punch biopsy of unaffected skin was taken as a negative control and preserved for RNA and protein. Sufficient tumor samples for both RNA and protein analyses on any given visit were obtained by three fine-needle aspiration biopsies taken from a palpable superficial lymph node: the same node was sampled over time for each canine. One FNA sample was placed immediately into Trizol to preserve RNA for subsequent RT-PCR analysis and in some cases microarray. The remaining two FNA samples were pooled and immediately frozen at −80°C for Western analysis. As for cell culture lysates, FNA sample protein concentrations were determined by Bradford assay.

### Microarray hybridization

Total RNA from tumor and skin samples was shipped on dry ice and sent to the Laboratory for Advanced Genome Analysis at the Vancouver Prostate Centre for microarray analysis (http://www.mafpc.ca/). Total RNA was used as a template to create labeled cDNA using MessageAmpTM Premier RNA Amplification Kit and MessageAmpTM III RNA Amplification Kit (Applied Biosystems) according to the manufacturer’s instructions. Labeled cDNA was hybridized to Agilent Canine Microarrays, which are comprised of more than 25,000 annotated genes. Scanning and data acquisition were obtained using the Illumina iScan scanner, raw data (idat files) were loaded into Illumina BeadStudio, without background subtraction, and exported for analysis. The data files have been deposited with the metadatabase Gene Expression Omnibus (GEO accession #GSE121242; https://www.ncbi.nlm.nih.gov/geo/query/acc.cgi?acc=GSE121242) as of Dec. 1, 2018. Sufficiently high quality of extracted RNA was not available for all samples or at all time points, limiting the overall assessment possible. We were only able to extract RNA from the skin control from canine 2. Canine 4 was unique in providing multiple quality protein and RNA samples over time as the canine progressed through clinical relapse and regression.

### Microarray data analysis

The microarray data were analyzed using the Limma R package. Raw probe intensity values were background-corrected using the backgroundCorrect function with method = “normexp” and quantile-normalized using the normalizeBetweenArrays function. X represented the 95^th^ percentile of the intensities of the negative control spots for a given array. We ignored any non-control probe for which there was no array having an intensity value for that probe that was more than 10% higher than X. The expression of a given gene was quantified by averaging the intensity values of all the probes corresponding to that gene. Finally, log2 fold-change values were calculated for each gene and for each treatment-control combination of interest. Venn diagrams were used to identify a common set of 290 genes that were expressed greater than 3 fold compared to skin controls in all four canines. Following identification of the 290 overexpressed genes that were shared between the four MDR canines, gene set enrichment analyses were conducted using Cytoscape 3.7.0 with ReactomeFIViz (2017 ReactomeFI Network Version). The 290 gene network was clustered to identify associated genes, and subsequent pathway enrichment was conducted. Pathways with a p-value and false discovery rate (FDR) of p<0.05 were included in results. The 290 gene list was also analyzed by STRING, version 10.5, as an independent method to identify clustered and associated genes. Analysis of over-represented transcription factor binding sites from the 290 up-regulated gene set was conducted using Cis-eLement OVERrepresentation (CLOVER; downloaded December 2018) with the transfac transcription factor motif database (2019). Analyses were run using a background sequence comprised of 1500 base pair sequences up-stream of transcription start sites in the genome. Sequences were retrieved from e!Ensembl database, Dog genes CanFam3.1. 1500 base pairs upstream of transcription start sites of each of the 290 genes were examined for the presence of transcription factor binding motifs, and motifs with a p-value < 0.05 (overrepresented) and p-value > 0.95 (underrepresented) identified.

## Author contributions

TGA was CoPI on the grant for the work, designed experiments, supervised the work, assisted in animal recruitment, interpretation of experiments, and writing of the manuscript. GFD performed the bulk of the cell culture experiments and Western assessment of animal tissues. LL performed the qRT-PCR experiments and provided cell culture expertise. BT and MW analyzed the microarray datasets. PF, DB and HM were undergraduate students who contributed to the work. ZEG performed the Cytoscape, CLOVER and transfact database analyses under CHE’s supervision. CHE performed the analysis required for GEO submission. FSV and FJV prepared the figure assessing APC substrate mRNA levels in patient samples. CG and VM-D were the veterinary oncologists responsible for animal recruitment, sample retrieval and animal care, and also assisted with study/experimental design, and were coinvestigators on the grant for the work. AK supervised the analysis of the microarray datasets. TAAH was the PI on the grant funding the work, designed the experiments, supervised the work, was involved in data interpretation and assessment, and writing of the manuscript.

## Acknowledgements

This work was supported by a grant to TAAH, TGA, VM-D, CG and AK from the Canadian Cancer Society (CCS) Innovation Grant program. CHE recieved support from the Natural Sciences and Engineering Research Council (NSERC) of Canada. FJV was supported by grants from the Canadian Institutes of Health Research (CIHR). ZEG received support from a CIHR Vanier PhD scholarship. MW was supported by an NSERC undergraduate summer scholarship, while PB and DB were supported by U of S College of Medicine Biomed summer awards.

---

**Supplemental Figure 1.**
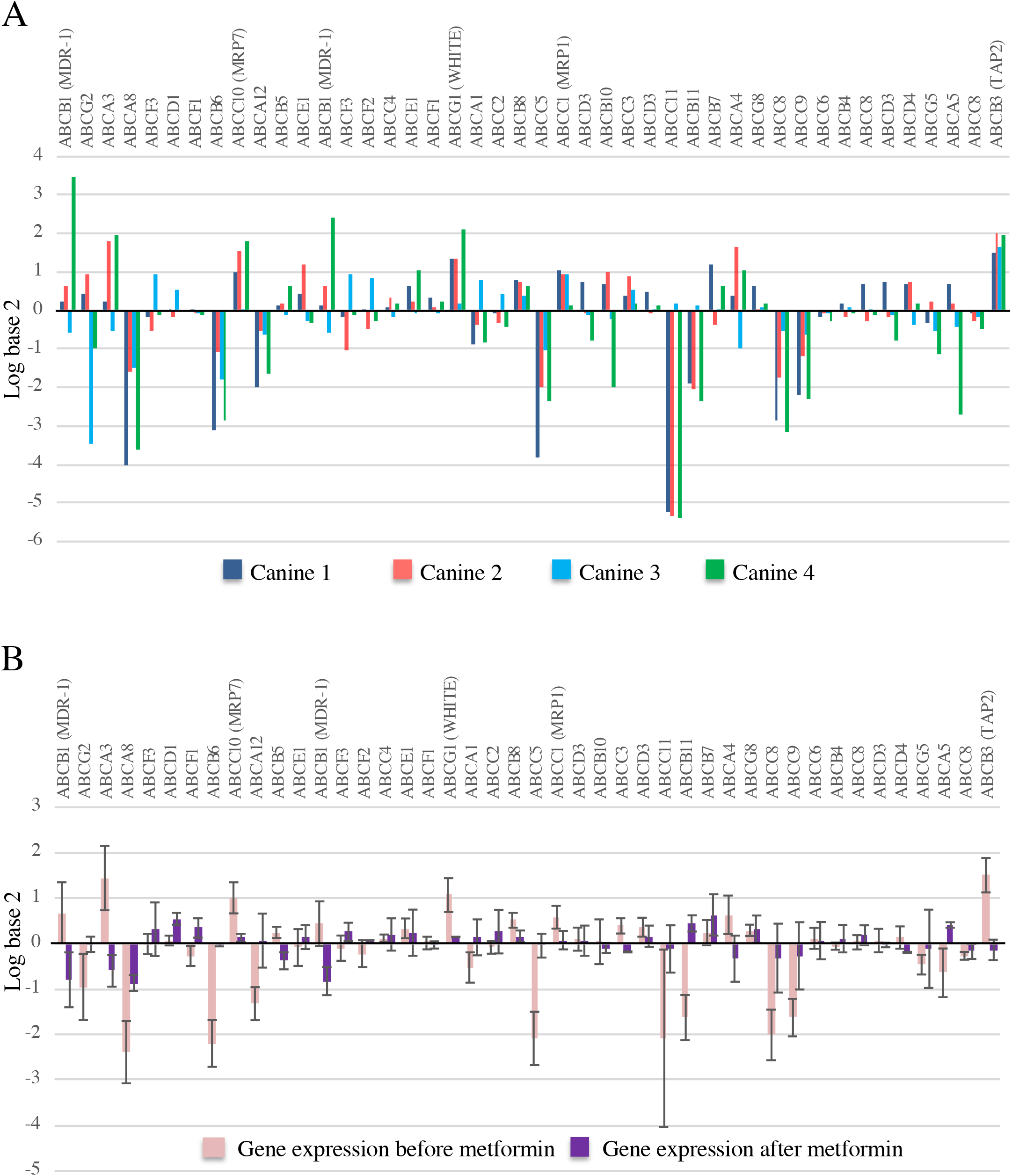
ABC transporters in general do not appear to play a critical role in the development of canine MDR lymphoma. **A.** Differential gene expression of the family of ABC transporters on the canine microarray was determined for each MDR canine sample. **B.** ABC transporter gene expression changes were averaged for the 2 canines (2 and 4) that were treated with metformin and analyzed by microarray. The light purple bars represent MDR tumor before metformin compared to controls. The bars in dark purple represent MDR tumors after metformin treatment compared with before treatment. The SEM was plotted.

**Supplemental Figure 2.**
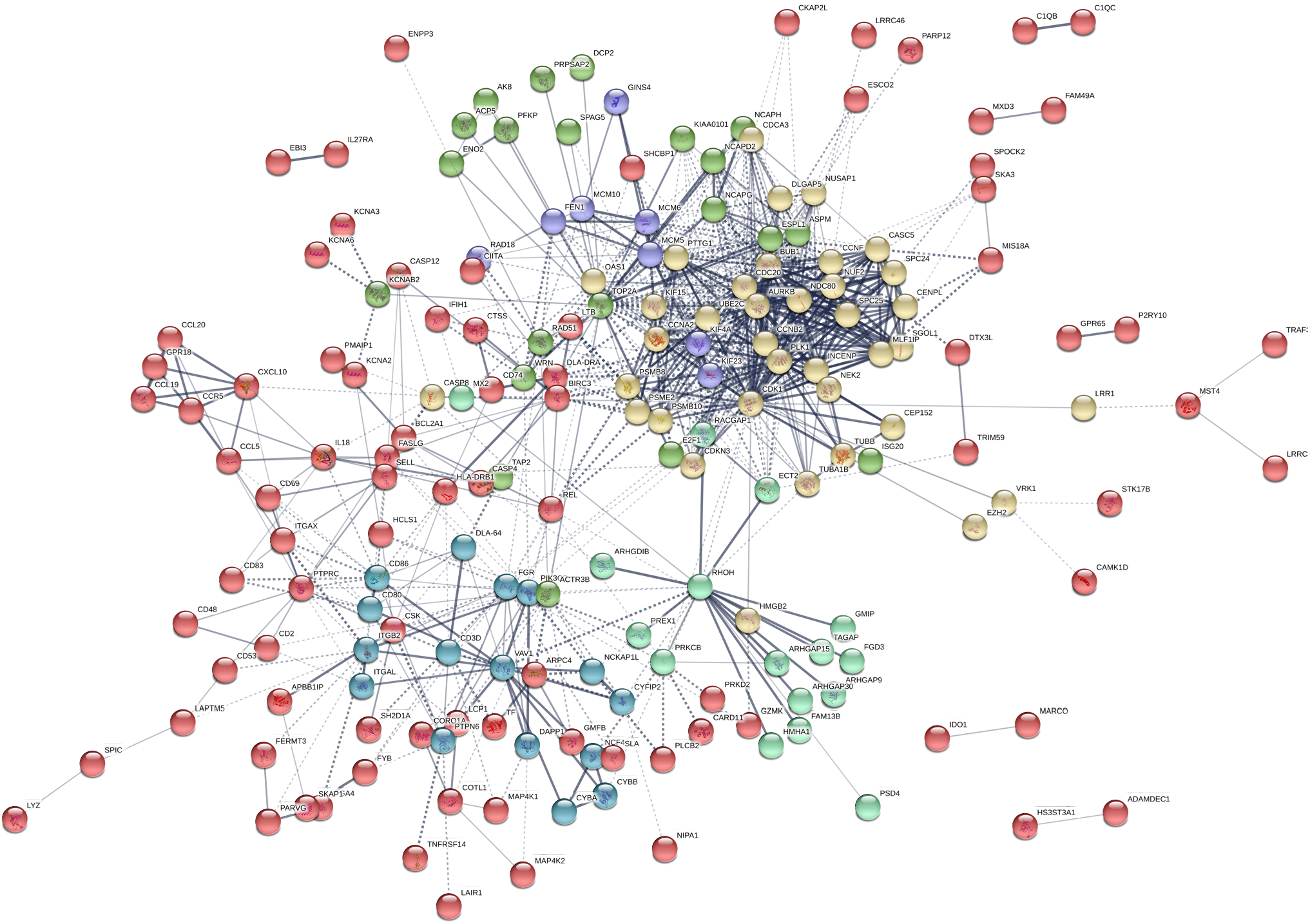
186 genes form the 290 gene set are predicted to form a highly interconnected network based on STRING. STRING analysis of the 290 common overexpressed genes within the tumors of MDR canines (Figure 2A) reveals a highly interconnected 186 gene set. A thicker connecting edge indicates higher confidence in the predicted interaction. The genes were grouped into 6 clusters, as shown by the different colored nodes. The yellow nodes are highly clustered and largely define genes involved in mitotic progression (see Supplemental Table 5). The green and purple nodes, also tightly connected to the yellow nodes, are primarily involved in chromosome maintenance and DNA repair. A significant number of these genes encode proteins known to be targeted by the Anaphase Promoting Complex (marked by an * in Supplemental Table 5). The majority of these genes are elevated in a variety of human cancer types.

**Supplemental Figure 3.**
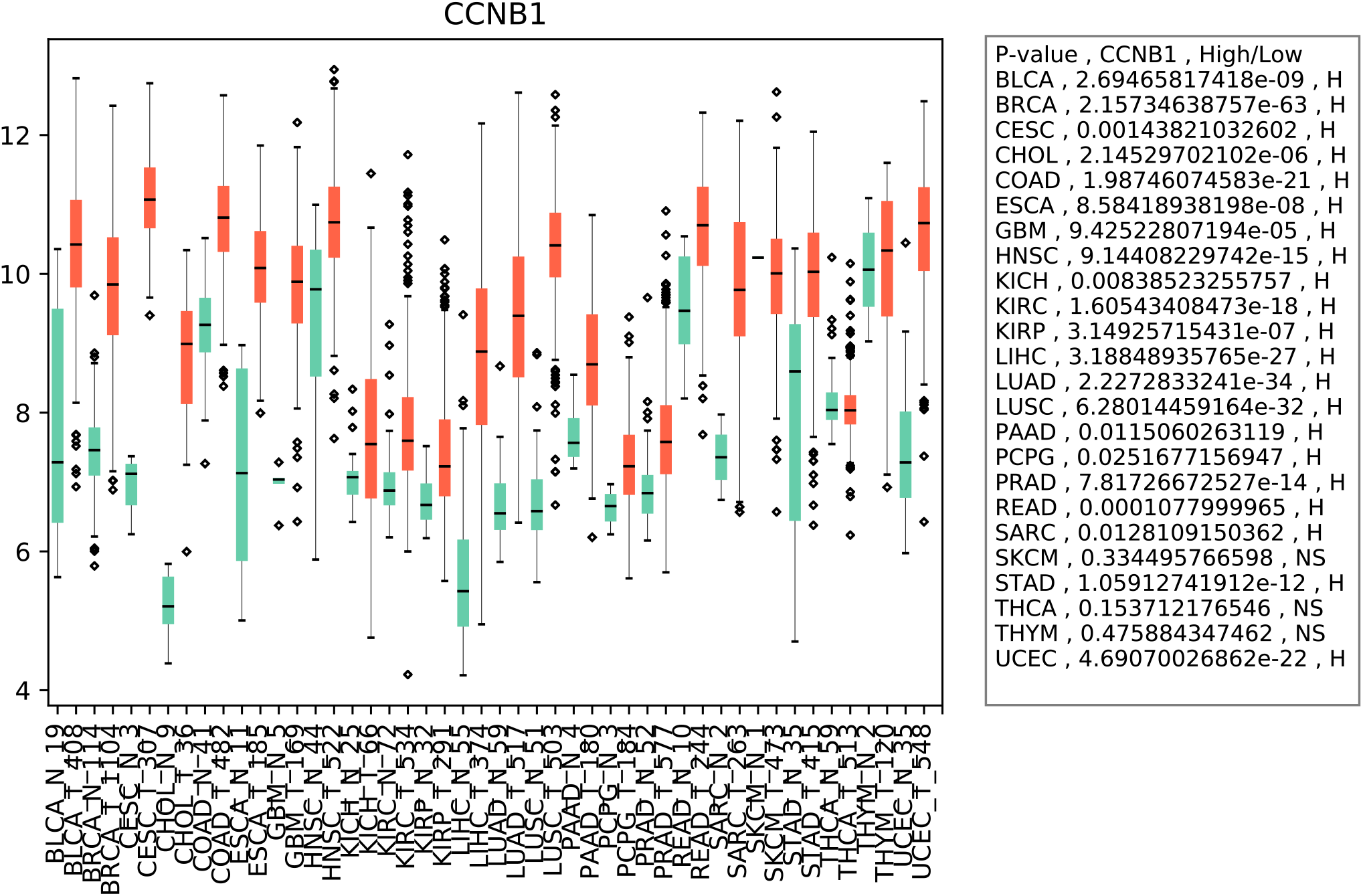
Expression scores for CCNB1 within 24 different types of cancer and normal tissue. Using the Cancer Genome Atlas (TCGA) (https://portal.gdc.cancer.gov/) database, we determined if expression of an APC substrate gene (CCNB1) in cancer patients is differentially regulated between normal and the tumor tissues. The numbers along the x-axis denote the number of patient samples in each cancer type. Statistical significance of the difference in expression between the normal and tumor samples are depicted for each cancer type. NS denotes not significant. The abbreviation of each cancer along the y-axis is represented as described in the TCGA portal.

**Supplemental Figure 4.**
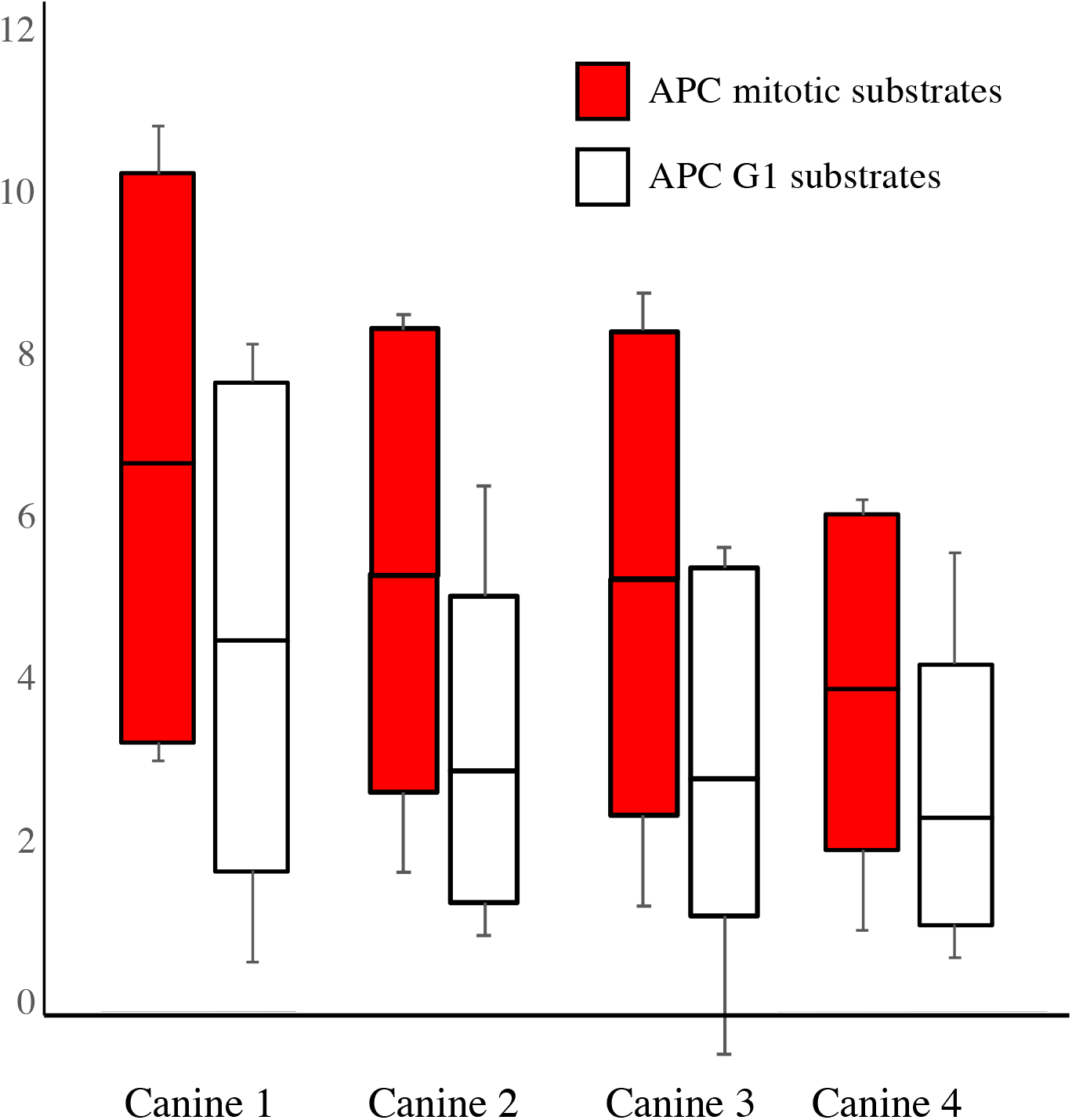
Expression of APC substrates is highly increased in canines with MDR lymphoma, with mitotic substrates showing a greater elevation than G1 substrates. The scores for all mitotic and G1 substrates from the 4 canine samples were averaged and plotted as shown. The top and bottom of the box define the third and first quartiles, respectively, while the median of the data divides the box. The whiskers define the error bars for the dataset.

**Supplemental Figure 5.**
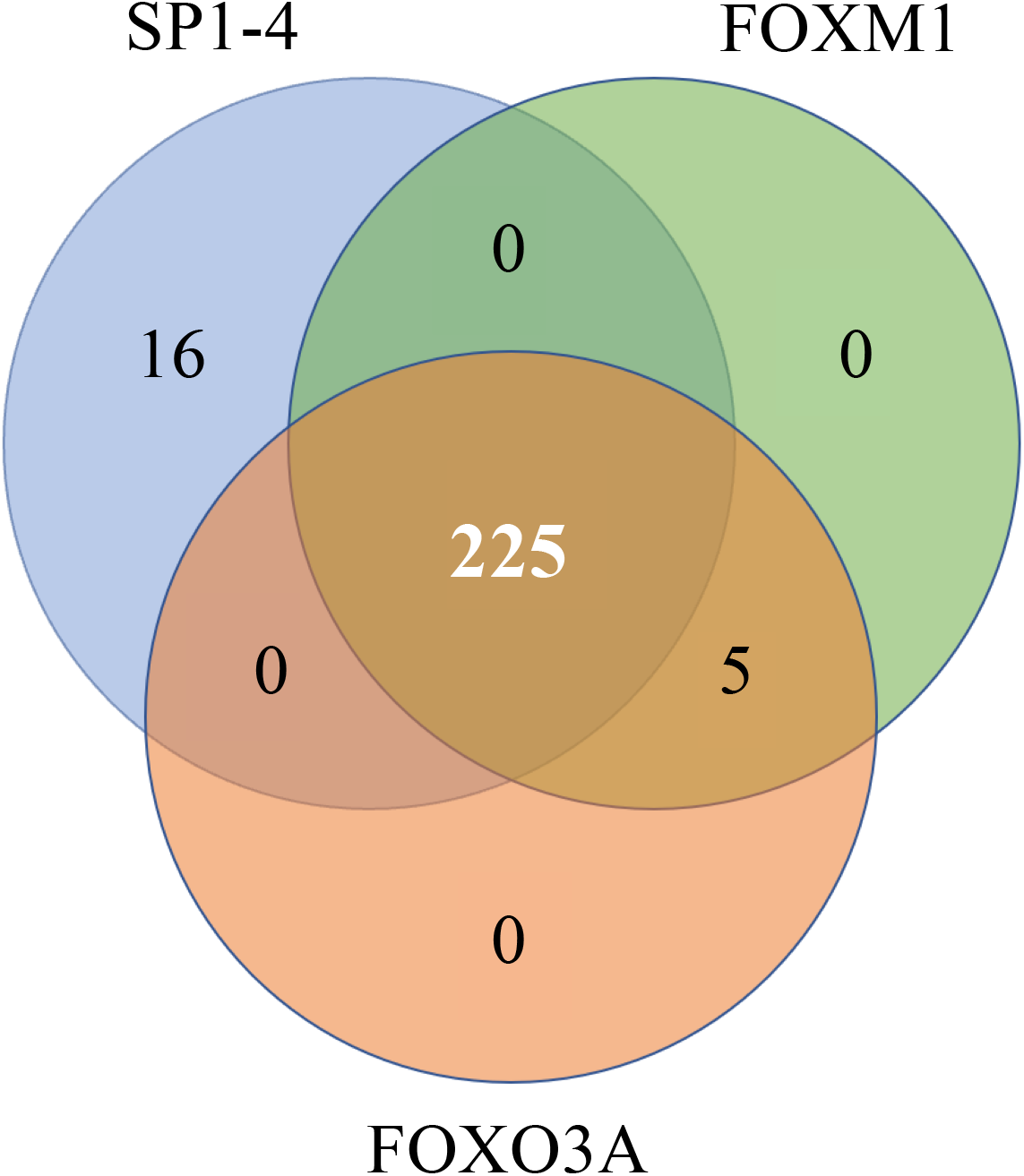
Binding sites within the promoters of the genes in the 290 gene set are enriched for recognition by FOXM1, FOXO3A and the SP1-4 class of transcription factors. A Clover analysis (https://www.ncbi.nlm.nih.gov/pubmed/14988425) using the Transfact database, identified enrichment for FOXM1 (230 genes), FOXO3A (230 genes) and SP1-4 (241 genes) promoter binding sites. A Venn diagram was used to identify overlaps between these datasets. 225 genes were found to contain binding sites within their promoters for all three sets of transcription factors.

**Supplemental Table 1.** Clinical characteristics of the MDR lymphoma canines used for the microarray analysis. All 4 MDR canines were male/neutered with recurrent B-cell lymphoma. All canines were clinically nonresponsive/MDR to therapy at the time of enrollment. Skin biopsy samples were taken prior to metformin treatment. Repeat FNA sampling was performed before and during MET addition to chemotherapy. MET: oral metformin in tablet form. OD: once daily. BID: twice daily. Stage III is generalized lymph node involvement of the front and back half of the body. CHOP: 4 drug chemotherapy cocktail given over 19 weeks including Cyclophosphamide, Doxorubicin (Adriamycin), Vincristine, Prednisone. CCNU is lomustine, an alkylating agent.

| Case ID number | Breed | Age | B vs T cell lymphoma | Stage | Chemotherapy at times of sampling | Clinical response to therapy | MET duration | MET dose | Time to death after MET | Overall survival (diagnosis to death) |
| --- | --- | --- | --- | --- | --- | --- | --- | --- | --- | --- |
| 1 | Golden Retriever | 10 yr | B | III | CHOP with MET | No, enlarged lymph nodes remained | 108 days | 250 mg OD | 184 days | 254 days |
| 2 | Lab Retriever mix | 5 yr | B | III | CHOP with MET | No, enlarged lymph nodes remained | 81 days | 500 mg BID | 87 days | 277 days |
| 3 | Retriever mix | 7 yr | B | III | DOX with MET | No, enlarged lymph nodes remained | 138 days | 500 mg BID | 143 days | 184 days |
| 4 | Lab Retriever mix | 11yr | B | III | CHOP failed then rescue with CCNU failed, then MET added to CHOP and remission | Yes/No, Remission attained for 8 weeks then relapse again | 84 days | 250 mg OD then 250 mg BID | 91 days | 261 days |

**Supplemental Table 2. Lists of genes upregulated >3 fold in the 4 MDR canines.** All genes differentially expressed >3 fold for the tumor samples, compared to control skin samples, were retreived from the microarrays. Venn analyses identified a common set of 290 genes overexpressed in all 4 MDR canines. STRING analyses identified a set of 186 genes within the 290 gene set that were highly connected.

**Supplemental Table 3. Cytoscape node list for the 146 gene set (see Figure 2B).** The 290 gene set was analyzed using the Cytoscape online tool. 13 interconnected nodes, composed of 146 genes, were identified that defined pathways involved in DNA repair and cell cycle progression through mitosis.

**Supplemental Table 4. Cytoscape network pathway enrichment for 146 gene set (see Figure 2B).** Network, Biological and Cell Component pathways were identified from the 146 gene network indicating the number of total genes in the complete gene set and the number found in our network. The enrichment was indicated by the p-value.

**Supplemental Table 5. Gene clustering from the STRING analysis identifies a set of genes within the 186 gene set that defines progression through mitosis as a key regulator of MDR development.** The clusters identified in Supplemental Figure 2 were listed and categorized in terms of function based on review of the literature. Genes highlighted in yellow define genes involved in mitotic progression, while genes highlighted in green define genes involved in DNA replication and repair. * denotes APC substrates; ** denotes kinetochore and Spindle Assembly Checkpoint associated proteins; *** denotes components of the chromosome condensin complex.

**Supplemental Table 6. Analyses of genes identified as responsive to clinical chemoresistance, remission, and relapse in canine 4.** A table comparing genes overexpressed >3 fold in the chemoresistant tumor, down-regulated >2 fold upon remission, and >3 fold following relapse in canine 4, identified 27 genes in common. STRING interactions were identified for each comparison. As above, genes highlighted in yellow define genes involved in mitotic progression, while genes highlighted in green define genes involved in DNA replication and repair. * denotes APC substrates; ** denotes kinetochore and Spindle Assembly Checkpoint associated proteins; *** denotes components of the chromosome condensin complex.

